# Sequence analysis allows functional annotation of tyrosine recombinases in prokaryotic genomes

**DOI:** 10.1101/542381

**Authors:** Georgy Smyshlyaev, Orsolya Barabas, Alex Bateman

## Abstract

**Background:** Tyrosine recombinases perform site-specific genetic recombination in bacteria and archaea. They safeguard genome integrity by resolving chromosome multimers, as well as mobilize transposons, phages and integrons, driving dissemination of genetic traits and antibiotic resistance. Despite their abundance and genetic impact, tyrosine recombinase diversity and evolution has not been thoroughly characterized, which greatly hampers their functional classification.

**Results:** Here, we conducted a comprehensive search and comparative analysis of diverse tyrosine recombinases from bacterial, archaeal and phage genomes. We characterized their major phylogenetic groups and show that recombinases of integrons and insertion sequences are closely related to the chromosomal Xer proteins, while integrases of integrative and conjugative elements (ICEs) and phages are more distant. We find that proteins in distinct phylogenetic groups share specific structural features and have characteristic taxonomic distribution. We further trace tyrosine recombinase evolution and propose that phage and ICE integrases originated by acquisition of an N-terminal arm-binding domain. Based on this phylogeny, we classify numerous known ICEs and predict new ones.

**Conclusions:** This work provides a new resource for comparative analysis and functional annotation of tyrosine recombinases. We reconstitute protein evolution and show that adaptation for a role in gene transfer involved acquisition of a specific protein domain, which allows precise regulation of excision and integration.

## BACKGROUND

Tyrosine recombinases (TRs) form a large family of proteins that perform site-specific DNA recombination in a wide variety of biological processes [1,2]. They are involved in post-replicative segregation of plasmids and circular chromosomes in bacteria, archaea and phages, protecting genome integrity upon cell division. The highly conserved Xer proteins (e.g. XerC and XerD in *E.coli*) resolve chromosome multimers formed after DNA replication in prokaryotes (reviewed in [3]), and the Cre recombinase separates dimers of the P1 phage genome (reviewed in [4]). Other TRs act as genetic switches, triggering phenotype variation within bacterial populations via DNA inversion or deletion [5–7]. In addition, TRs drive the mobilization of ‘selfish’ genetic elements, including various phages and transposons. Some mobile elements hijack host-encoded Xer proteins [8,9], while others encode distinct TRs to promote their own integration and transfer in bacterial genomes. Prominent examples of TR-carrying mobile elements are the integrative and conjugative elements (ICEs), also referred to as conjugative transposons. ICEs combine features of phages and plasmids, because they can both integrate into genomes and disseminate by conjugative transfer [10]. A large number of ICEs and related non-autonomous mobilizable elements are present in diverse bacterial taxa and provide efficient vehicles for horizontal transfer of genetic traits such as virulence and antibiotic resistance genes [11–15]. TRs are also found in antibiotic resistance-carrying non-conjugative transposons [16,17], and they are essential for the activity of integrons that capture antibiotic resistance genes into complex functional units (reviewed in [18]). Thus, TRs contribute to the emergence and spread of bacterial virulence and antibiotic resistance on a multitude of ways and charting their distribution and specific functions is instrumental to understanding genetic transfer traits. Yet, functional annotation of TRs in genome sequence data has been challenged by their large diversity and poor phylogenetic characterization, leaving the vast majority of TRs without a reliable functional prediction. Previous sequence analysis of a limited number of well-annotated TRs suggested that sequence features may help predict specific function [19,20]. However, a comprehensive classification of TRs is still missing and the diversity, phylogenetic relationships and specific sequence features of different TR groups are poorly understood This compromises our understanding of site-specific recombination mechanisms, hampers automated identification and functional annotation of TRs and TR-carrying genetic elements, and ultimately limits our knowledge of microbial adaptation and gene transfer pathways. In the past decade, the number of available genome sequences has multiplied, opening new possibilities for studying TR diversity and evolution.

Here, we have used a comprehensive search protocol to assemble a complete set of bacterial and archaeal Xer-related TRs from protein databases. By sequence analysis and classification, we show that integrases from ICEs and phages emerged from a Xer-like ancestor by acquisition of a so-called arm-binding (AB) domain. In turn, phylogenetically distinct ‘arm-less’ TRs function as chromosome dimer resolution proteins, integron integrases or transposases/resolvases of non-conjugative elements. TRs within phylogenetic groups share specific features, such as preferred host taxonomy and characteristic structural perturbations. Based on this TR classification system we further identify and characterize new ICEs, demonstrating the power of our sequence based functional annotation. These results provide insight into the diversity, distribution and evolution of this abundant protein family and create a resource for annotation of novel TRs and mobile elements.

## RESULTS

### Identification and analysis of tyrosine recombinases

Previous structural and sequence analysis indicated that most TRs have two functional domains: the core-binding domain (CB) binds the core DNA site, and the catalytic domain (CAT) catalyzes DNA cleavage and joining reactions for recombination [21–23]. Some TRs have an additional N-terminal AB that recognizes so-called arm DNA sites. Crystal structures indicate that CAT has an overall similar fold in diverse TRs [24–27] and comparative sequence analyses from a subset of CAT sequences revealed two highly conserved regions (referred to as boxes) and three patches with less significant conservation [22,23]. These conserved positions correspond to the catalytic residues (i.e. the tyrosine nucleophile and the catalytic pentad RKHRH [1]) and the hydrophobic protein core. The CB domain is much less conserved on the sequence level, but its structure generally features a bundle of four alpha-helices [21]. Finally, the AB domain is the most variable with remarkable structural and sequence diversity between TR family members [28–30].

To comprehensively analyze the diversity of TRs we employed the following strategy: First, we performed an iterative jackhammer search against the UniProt reference proteomes database using the XerD protein from *Escherichia coli* as a query. The resulting sequences were then made non-redundant, aligned, and the phylogenetic tree was reconstructed using the PhyML package (Figure 1A, See Materials and Methods for more details). Based on the phylogeny, TRs were then divided into subgroups, for which we then created HMM profiles. These profiles were then used for a subgroup-specific search in UniProt reference proteomes. For the resulting sequences, we then created sequence logos, which showed the conserved regions within subgroups and the specific differences between subgroups (Figures 2-5). Finally, we mapped all TR proteins to their genomes of origin, and plotted the distribution of each subgroup among taxa (Figure 1B). We also extracted the fifty most abundant TR proteins and charted their distribution, classification and putative function (Figure 1C).

**Figure 1.**
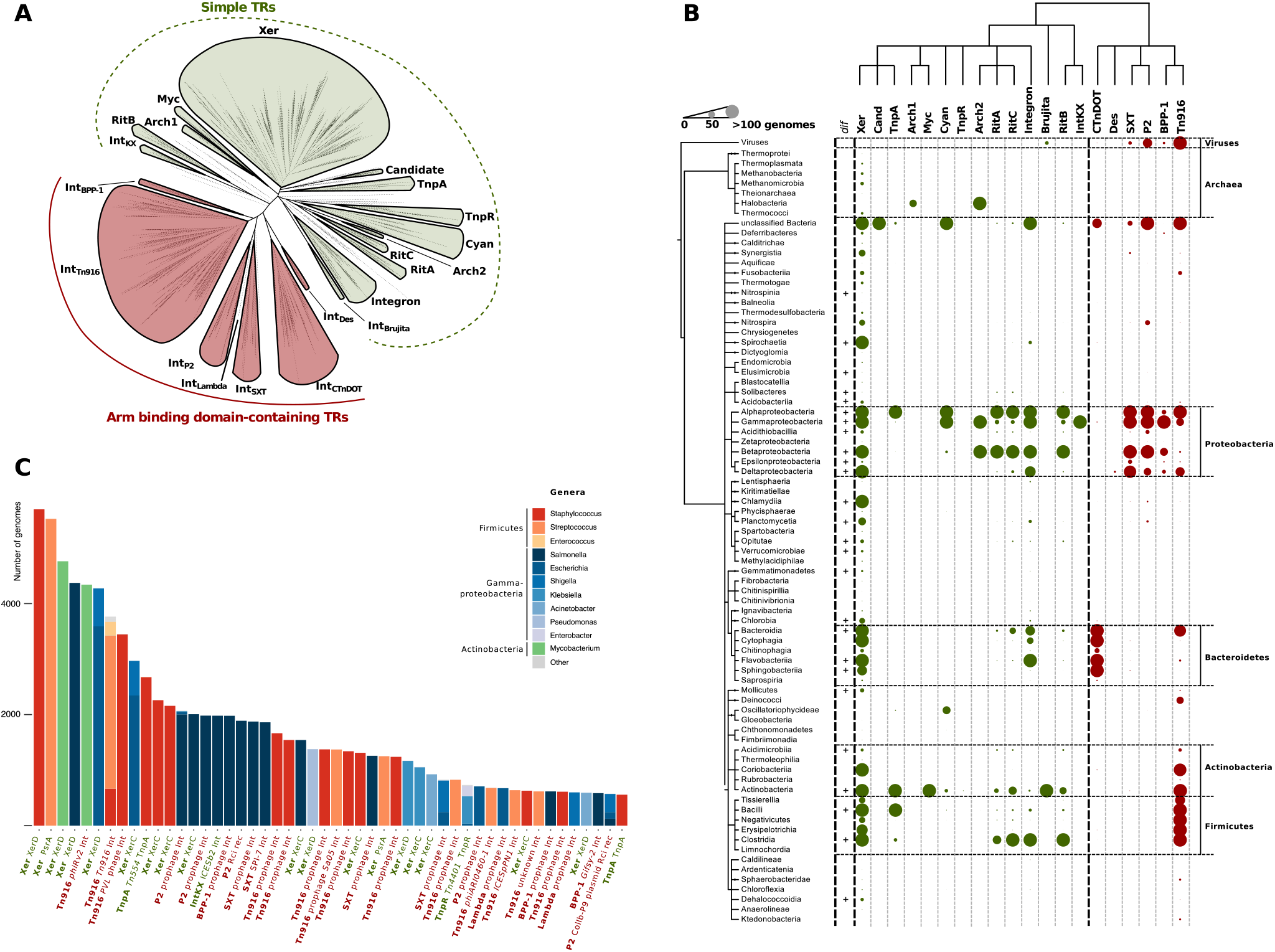
**A.** Maximum likelihood phylogenetic tree of tyrosine recombinases. Two major groups of TRs: simple and AB-containing TRs are highlighted in red, and green, respectively. Subgroups of the TRs are shown as leaves in the tree. Statistical support was evaluated by aBayes and for all of the subgroups its value is more than 0.98. **B.** Taxonomic distribution of tyrosine recombinases. At the top a schematic tree of the tyrosine recombinase phylogeny corresponding to the panel A is shown (only nodes with statistical support of more than 0.98 are shown). Phylogeny of the bacterial taxa is shown on the left. The abundance of TRs belonging to a specific TR subgroup in a particular taxon is indicated by different size dots in the plot. The exact number of genomes are available in the Supplementary Table 2. **C.** The fifty most abundant tyrosine recombinase proteins found in the genomic sequences available from NCBI. The bars indicate TR abundance in different bacterial taxa with distinct colors. NCBI GI numbers for all the sequences are available in the Supplementary Table 3.

**Figure 2.**
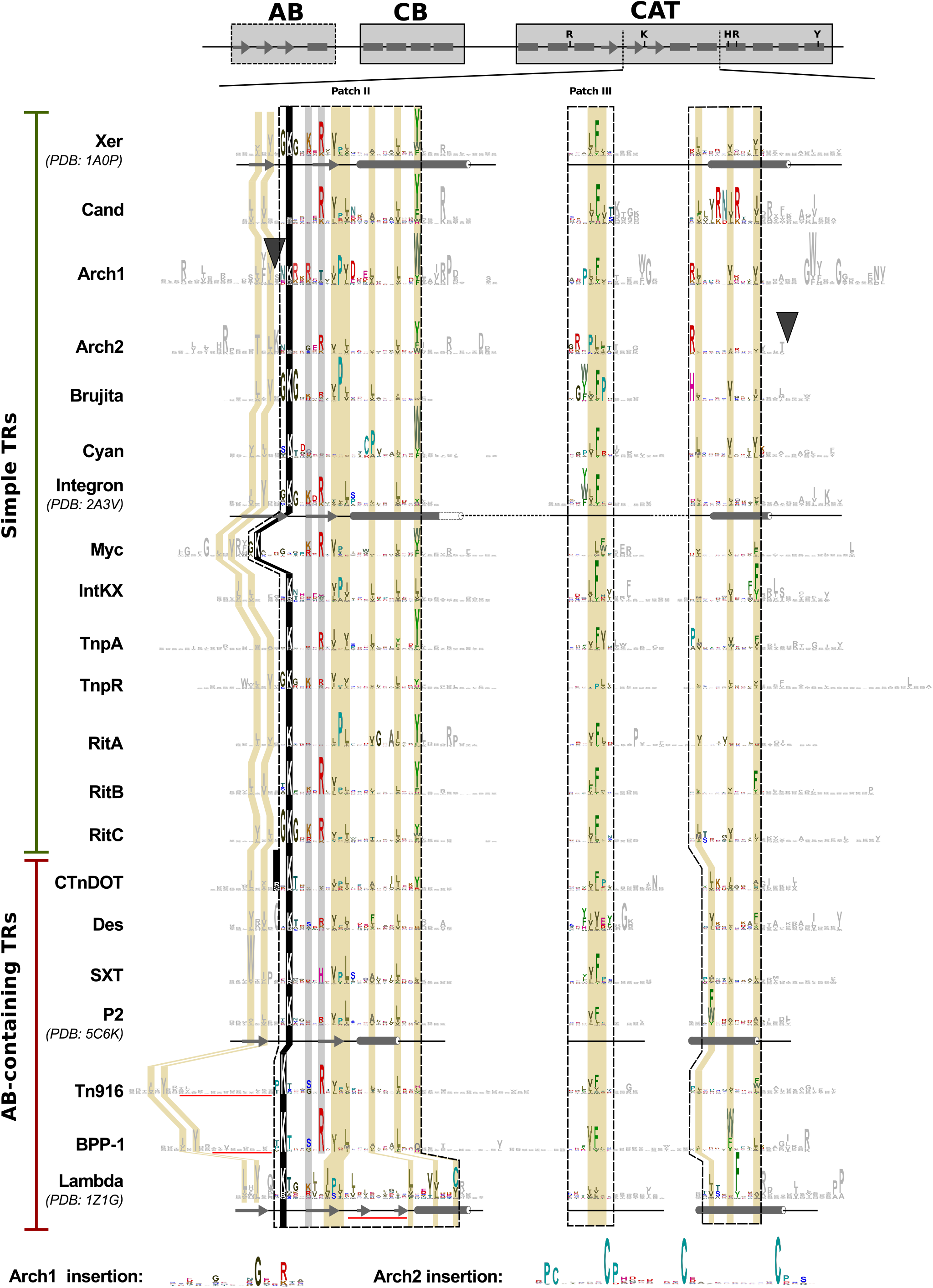
Conservation analysis of the catalytic domain (CAT) of tyrosine recombinases (N-terminal part). Manually aligned web logos produced after hmmsearch analysis against the UniProt reference proteomes database for each of the subgroups. Conserved boxes and patches are manually aligned and highlighted with color corresponding to different residues. The tyrosine nucleophile and the catalytic RKHRH pentad are highlighted with black background. Positions corresponding to conserved residues located near the active site pocket are highlighted in grey. The conserved hydrophobic residues that form the core of the catalytic domain are highlighted in light brown. The secondary structures of a representative subgroup member were retrieved from PDB accessions, where available, and are shown below the logos.

**Figure 3.**
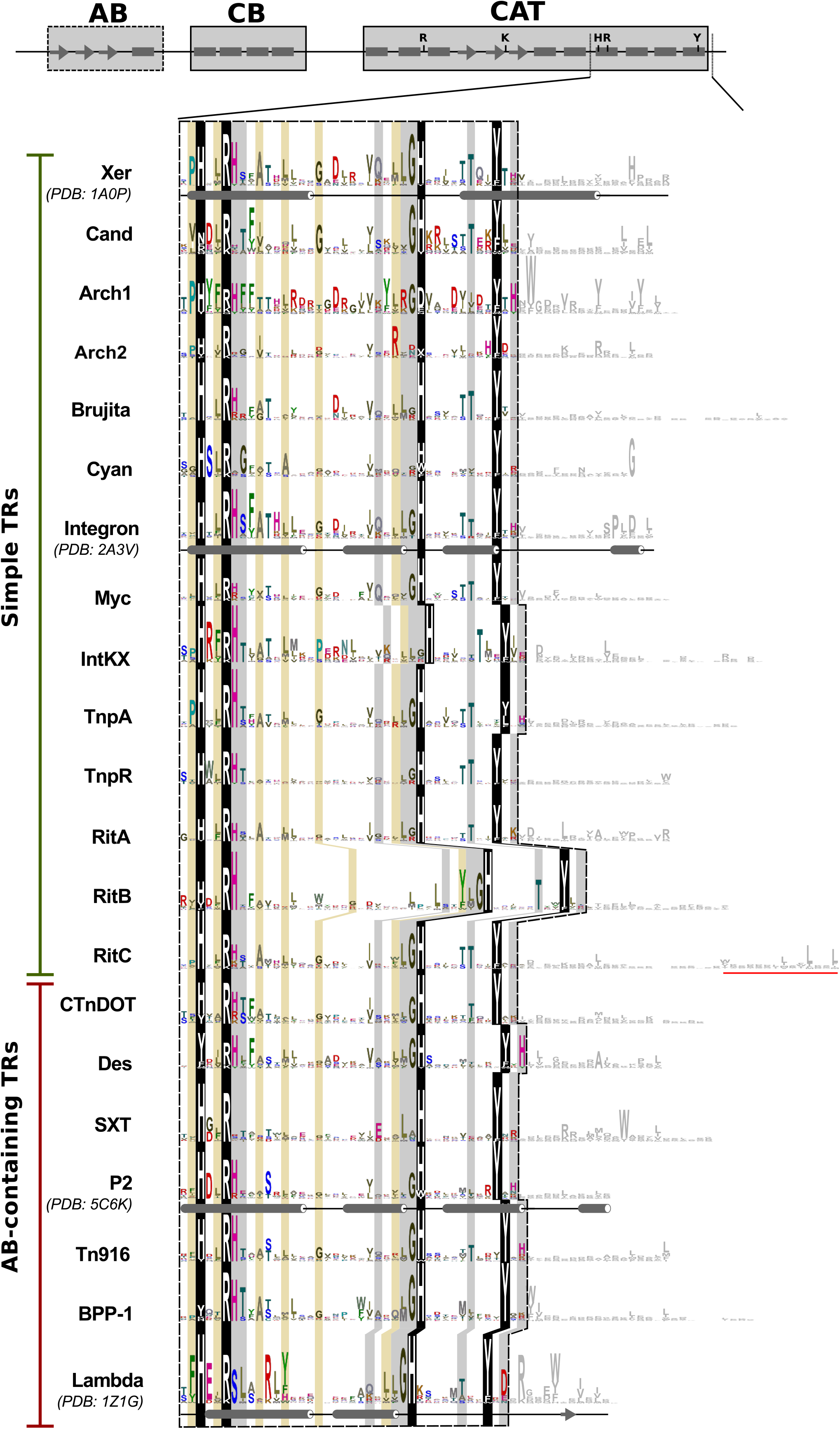
Conservation analysis of the catalytic core domain of tyrosine recombinases (central part). Manually aligned web logos produced after hmmsearch analysis. Color code as in Figure 2. The position of insertions in the Arch1 and Arch2 TRs are indicated with black wedges in the corresponding logos and are shown in fully at the bottom of the figure. Insertions in BPP-1, Tn916 and λ TRs are underlined in red.

**Figure 4.**
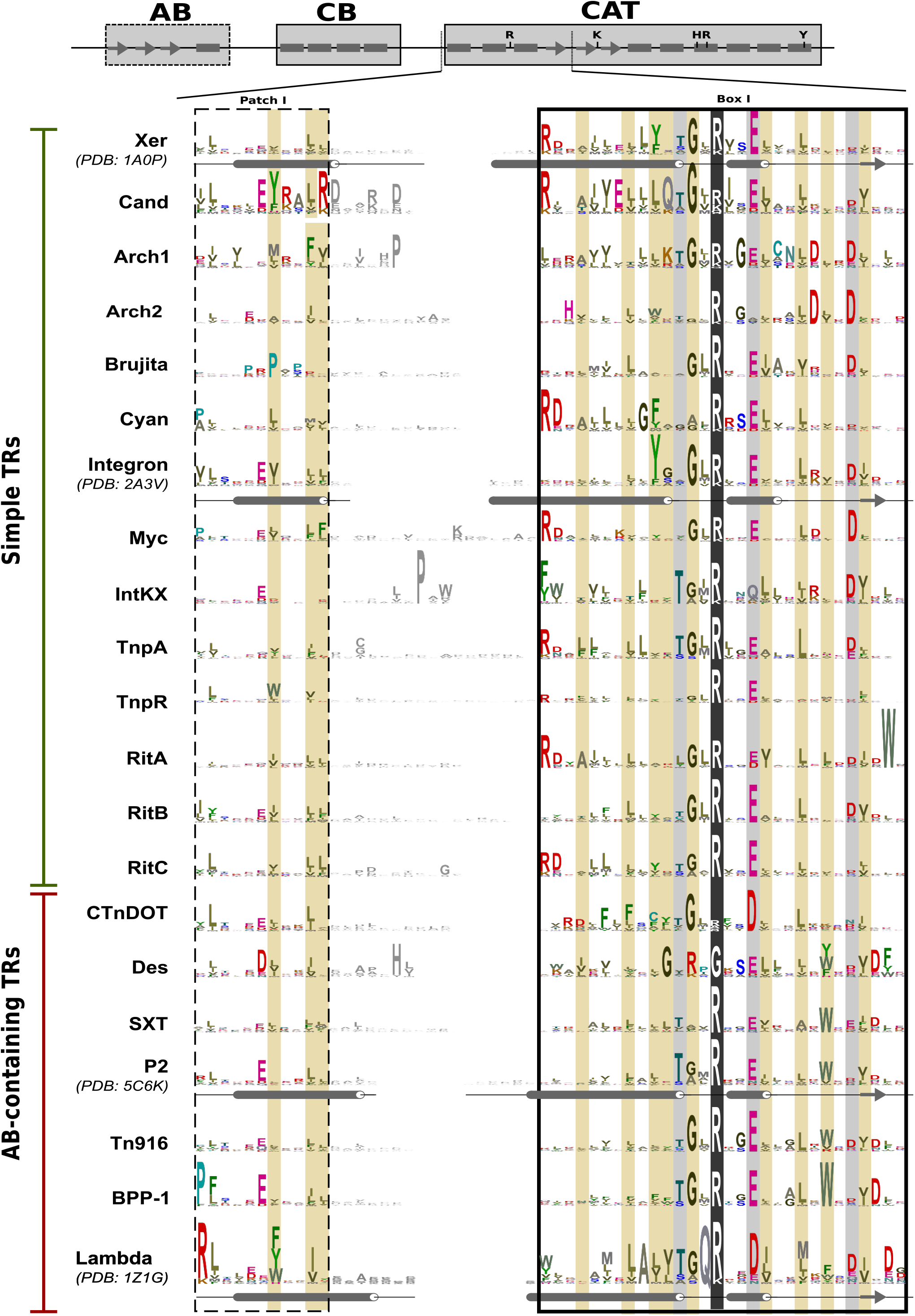
Conservation analysis of the catalytic core domain of tyrosine recombinases (C-terminal part) as in Figures 2 and 3. Insertions in RitC TRs are underlined in red.

**Figure 5.**
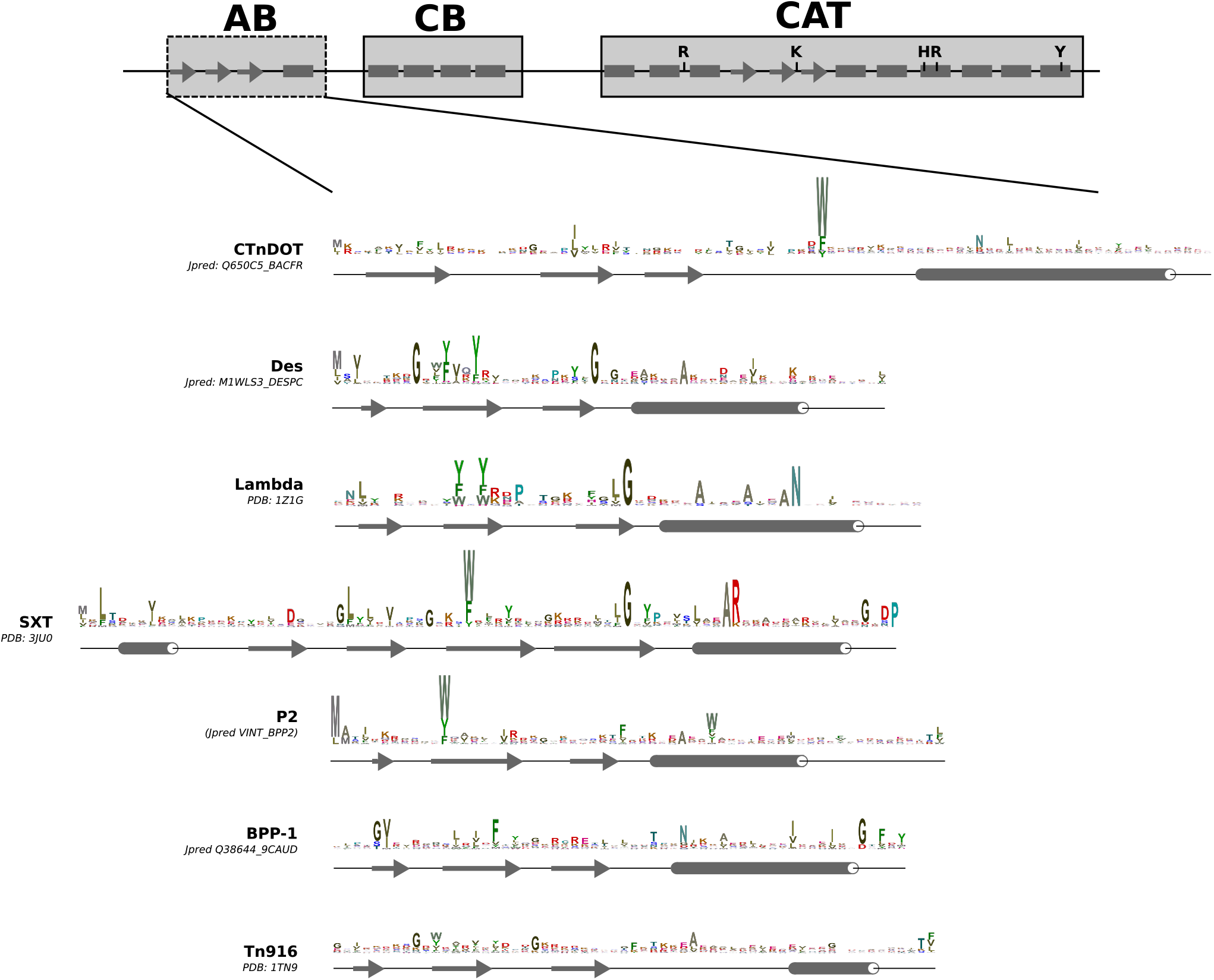
Conservation of the arm-binding domains of tyrosine recombinases. Web logos were produced after hmmsearch analysis against UniProt reference proteomes database for each of the subgroups. For each of the subgroups secondary structures were predicted using Jpred or retrieved from corresponding PDB accessions and are shown below the logos.

This analysis showed that all TRs can be classified in two major phylogenetic groups: simple ‘arm-less’ TRs, and AB-containing TRs (Figure 1A). As reflected in their names, the major difference between these two groups is the presence or absence of an AB domain. Within these main groups, smaller subgroups were identified (Figure S1), which are distinguished by specific structural and sequence features within a generally conserved domain architecture. taxonomic distribution. Remarkably, TRs within subgroups have a characteristic taxonomic distribution and specific functions highlighting the applicability of the sequence analysis for functional annotation. In the following sections, we first describe characteristic features of each major subgroup and then focus on using the classification system to identify new ICEs in bacterial genomes.

### Simple TRs

Simple TRs usually contain two domains – CB and CAT (Figures 1A and S1)The group can be further classified into fourteen subgroups.

The largest subgroup, Xer includes proteins responsible for chromosome dimer resolution in bacteria and archaea, such as XerC/D, XerH, XerS and XerA [31–34]. In addition, an integrase of the Helicobacter conjugative transposon TnPZ (XerT) [35] and a highly abundant recombinase that drives production of different methyltransferase gene alleles in *Streptococcus pneumonia* [5,6] (PsrA; Figure 1C). As judged by the sequence logos, Xer is the most evolutionary conserved subgroup: it has highly conserved residues also outside of the active site pocket and the hydrophobic core of the catalytic domain. Xer is also the most widely distributed subgroup. It is present in all bacterial and archaeal classes where TRs were found, with the exception of Oscillatoriophycideae, Ktedonobacteria, Rubrobacteria, Deinococci and Halobacteria (Figure 1B, Supplementary table 2). Such broad distribution is consistent with the essential role of Xer proteins in chromosome dimer resolution in bacteria and archaea. In the remaining taxa, other class-specific simple TRs may complement Xer function. For example, in Halobacteria two specific types of simple TRs were found, named here Arch1 and Arch2, that contain a distinct insertion discriminating them from other TRs (Figure 3). Similarly, Oscillatoriophycideae contain members of a distinct simple TRs subgroup, named here Cyan (after Cyanobacteria – a phylum of the class). Another subgroup of simple TRs, Cand is closely related to Xer, and its members were found exclusively in unclassified ‘Candidate’ bacterial phyla that was recently recognized as a ‘microbial dark matter’ [36]. Finally, the Myc subgroup unites proteins with unknown function that are found only in Actinobacteria and cluster together with Arch1 representatives.

In addition to putative chromosome dimer resolution proteins, the simple TRs also include multiple subgroups related to mobile elements. First, TRs of the RitA, RitB and RitC subgroups are part of the three TRs of the so-called ‘recombinases in trio’ transposable elements [37]. In addition, the RitB subgroup includes a putative integrase of ICEBmu17616-1 conjugative transposon from *Burkholderia multivorans* (Supplementary Table 1). Notably, our conservation analysis suggests that TRs from the RitA clades have two sequential CB domains (Figure S3). On the other hand, RitC TRs seem to have an additional C-terminal conserved helix (as predicted by Jpred), which is absent in all other TRs (Figures 4 and S3). The TnpA subgroup is formed by TRs related to the TnpA transposase from transposon Tn554 [38]. Members of this subgroup have several additional segments that are missing in all other TRs, such as a beta strand insertion in the CAT, an alpha helix insertion between CB and CAT and an N-terminal insertion of two beta-strands (Figure S3). The Integron-like subgroup is marked by the well-studied Integron integrase, IntI, that is responsible for capturing gene cassettes by the integron recombination platform [39]. In addition, this group includes TRs found in IS91 and IS3 transposons [40]. IntI contains an insertion of several alpha helices within the canonical CAT domain fold that is needed for specific DNA recognition [41]. However, this seems to be a feature specific to IntI only, as it is absent in the closest homologue TRs from IS91 and IS3 and in the conservation logo of the subgroup in general (Figure 3). TRs from the TnpR subgroup include the TnpR protein from the carbapenemase-carrying Tn4401 [42] and related proteins found in beta-and gammaproteobacteria. TnpR is one of the most abundant TRs in bacterial genomes and can be found in diverse gammaproteobacteria, suggesting that Tn4401 may be highly abundant in this bacterial class (Figure 1C). Interestingly, many of the proteins belonging to the TnpR subgroup have a helical N-terminal domain DUF3701 (PF12482, Figure S1). The function of this domain is unknown, but it clearly contains a HHH motif, suggesting DNA binding. The Int_Brujita_ subgroup is formed by TRs of mycobacterial phages (Figure 4), including the Brujita phage TR protein, which was recently shown to performs integration of the phage into the bacterial genome [43]. Finally, the Int_KX_ subgroup (defined in [44]), comprises closely related TRs of ICEs (such as PAPI-1, ICESb2 and ICEEc2) that are known pathogenicity islands found in human pathogenic gammaproteobacteria [44–49]. These latter examples indicate that simple TRs can perform integration/excision of mobiles elements, like ICEs and phages.

### Arm-binding domain-containing tyrosine recombinases

The documented examples of these TRs are found mostly as integrases of phages/prophages or ICEs. Their main feature is the presence of an arm-binding domain, although few members seem to lack it (Figure S1). The AB-containing TR group consists of six subgroups; below we discuss the characteristic features of all subgroups.

### Int_CTnDOT_ subgroup

The Int_CTnDOT_ subgroup includes integrases of the ICEs CTnDOT and NBU1 [50,51], as well as related TRs found predominantly in Bacteroidetes. In addition, some Int_CTnDOT_ subgroup members that are related to the integrases of the Salmonella genomic island 1 (SG1) – a mobilizable element containing an antibiotic resistance gene cluster [52,53] – were also identified in gammaproteobacteria (Figure 1B).

Regarding domain composition, initial Pfam annotation suggested that, Int_CTnDOT_ TRs contain two domains – CB and CAT, similar to the simple TRs. However, the CB domain, was predicted to belong to a different domain family (SAM5, Pfam family: PF13102) and was longer than that of the simple TRs. Subsequent conservation analysis revealed that the N-terminal part of Int_CTnDOT_ TRs can be split into two domains: a canonical CB and an N-terminal AB (Figure S1, the AB is shown in Figure 5).

Furthermore the integrase of the CTnDOT element was previously predicted to have an AB that is composed of three beta strands and one alpha helix [54], which is similar to the secondary structure composition of the arm-binding domain of λ integrase, and the corresponding arm DNA sites were also identified [55,56]. Since the majority of the Int_CTnDOT_ subgroup members were predicted to have SAM5 type N-terminus, this bipartite structure seems to be conserved within the subgroup. Therefore, we have split the SAM5 domain into two domains: an Arm-binding domain (Arm-DNA-bind_5 with new Pfam accession PF17293) and SAM5 (Figure S1). This change is now available in the new version of Pfam (Pfam 31.0).

In the CAT domain, TRs of Int_CTnDOT_ subgroup feature a weaker conservation of the first arginine in the RKHRY pentad (conserved R, Box I, Figure 2). An arginine at this position is present in the integrases of the non-autonomous NBU1 and NBU2 elements, and of Tn4555, but is absent from the integrases of CTnDOT, ERL (S) and Tn5520 elements [57]. In the CTnDOT integrase, this arginine was proposed to be functionally substituted by a different arginine located at the end of first beta-strand [58]. We find that this alternative arginine is indeed conserved in many integrases in the Int_CTnDOT_ subgroup (see conserved R in Int_CTnDOT_ logo in Figure 3). This indicates that the TRs of this subgroup may have the catalytic arginine either in the canonical location (NBU1), or in the alternative location (CTnDOT), resulting in an apparent weaker overall conservation of both positions in the entire subgroup.

### Int_Des_ subgroup

Int_Des_ is a small group of AB-carrying TRs. Its members are found only in the genus Desulfovibrio of Deltaproteobacteria (Figure 1B). This subgroup features specific perturbations in the catalytic core: namely the first arginine of the RKHRH pentad is substituted to glycine and the first histidine is substituted to tyrosine (Figures 2 and 4). The role of these TRs remains unknown.

### Int_SXT_ subgroup

Int_SXT_ subgroup comprises integrases of several ICEs, genomic islands and phages and related TRs. Our analysis revealed that a characteristic feature of this subgroup is the presence of an N-terminal DUF4102 domain (Figure S1). This was previously annotated as an AB of genomic island integrases [30] and contains an additional N-terminal beta strand and an alpha helix compared to ABs of other TRs (Figure 5). Further phylogenetic analysis revealed that there are several lineages within the Int_SXT_ TRs (Figure S4). Integrases of P4 and Sf6 phages, which insert into tRNA genes, cluster together with TRs of ICEs that also target tRNA genes (P4 lineage, Figure S4) [19,59]. Similarly, integrases of epsilon15 phage, the CMGI-3 element from *Cupriavidus metallidurans* and integrases of related elements form a separate lineage and all target the *guaA* gene [60,61] (epsilon15 lineage, Figure S4). A similar pattern is seen for integrases of the Enterobacterial cdt1 phage, the SXT element and closely related ICEs, all of which target the *prfC* gene [62,63] (SXT lineage, Figure S4). Thus, all members of each Int_SXT_ lineage integrate their diverse mobile elements into specific locations, probably due to characteristic features in the integrase sequences that define their target site choice. In addition, the presence of both ICE and phage integrases in most lineages suggests that Int_SXT_ TRs of ICEs and phages diversified multiple times during their evolution or that phages and ICEs frequently interchange their integrases. The latter is also supported by the recent study suggesting that ICEs with different conjugation machineries may have very closely related integrases [64].

### Int_P2_ subgroup

The Int_P2_ subgroup contains the integrases of some proteobacterial phages, such as HP1 and P2. Crystal structures of the CAT domains from these phage integrases have previously revealed high similarity to other CAT structures [26,65]. The AB domains in this subgroup resemble the classical λ -like AB, with three beta strands and one alpha helix (Figure 5). Another interesting member of this family is the Rci recombinase, which regulates R64 plasmid conjugation by enabling the generation of diverse pili proteins [66–68]. The Rci recombinase fulfills a similar function as some of the phylogenetically unrelated simple Xer TR plays in phase variation, perhaps showcasing an example of convergent evolution. Although our and other [59] phylogenetic reconstructions suggest that Int_P2_TRs may be related to λ-like integrases, this clustering is not well supported (Figures 1A and S1). λ integrase is one of the best described tyrosine recombinases [69] that is often used as a prototype for the superfamily. However, our analysis rather indicates that it is quite different from other TRs [22]. Major differences in the CAT domain include an insertion of two beta strands after the third core beta strand and the replacement of the final C-terminal alpha helix with a beta strand (Figures 3 and 4).

### Int_BPP-1_ subgroup

The main distinguishing feature of the Int_BPP-1_ subgroups is the presence of a beta-strand insertion between the second and third beta strands in the canonical CAT domain fold (Figure 3). Members of this subgroup are found in gamma-and betaprotobacteria and phages (Figure 1B). Examples include putative integrases of the Bordetella BPP-1 phage, the Stx2a phage and the Salmonella Gifsy-2 phage [70–72], the latter being the most abundant protein of the subgroup (Figure 1C). Int_BPP-1_ TRs feature a distinct AB, DUF3596 (Pfam family: PF12167, Figure S1), which however exhibits a canonical three-beta-strand/one-helix structure (Figure 5). Members of the family also have relaxed conservation of the last histidine in the RKHRH pentad of the catalytic core (Figure 4).

### Int_Tn916_ subgroup

As Int_BPP-1_, the Int_Tn916_ subgroup also features a beta-strand insertion between the second and third beta strands in CAT (Figure 3). Recent structural and biochemical work on the Tn*1549* integrase from this subgroup has shown that the beta-insertion is essential for shaping the DNA substrate for recombination [73]. The N-terminal region was in some cases predicted as Integrase_AP2 domain by Pfam (Pfam family: PF14657), but in many TRs no Pfam domains could be detected (Figure S1). However, our additional conservation analysis revealed that members of the subgroup contain a conserved AB with the canonical three-beta-strand/one-helix structure (Figure 5), as seen in the Tn916 Int AB structure [74]. Int_Tn916_ subgroup members are most highly represented in phages and gram-positive bacteria, but are also found in other taxa, such as Fusobacteria, Synergista and Chlamydia (Figure 1B). The most abundant TRs in this subgroup are the Mycobacterium phiRV2 prophage integrase [75], found in the genomes of more than 4000 Mycobacterium strains, and the integrase of the tetracycline resistance-carrying Tn916 element [11] (Figure 1C). As in the Int_SXT_ subgroup, the phage and ICE TRs do not generally form separate clusters. Instead, most lineages within this subgroup contain integrases from both ICEs and phages. For example, many actinomycete ICE TRs (previously referred to as AICEs [76]) cluster into a single lineage with the integrases from actinobacterial phages, representatives of which seem to target tRNA genes (pSAM2 lineage, Figure S5). Interestingly, in contrast to most other TRs, those from the Tn916 lineage, including Tn*1549* integrase, have relaxed insertion site preferences and integrate their ICEs into AT-rich regions without a strict sequence specificity [77–79]. This feature might have contributed to the success of the Tn916 element in the genomes of gram-positive bacteria.

### Identification and classification of integrative and conjugative elements

The vast majority of the TRs have no annotation of their putative function in the protein or genomic databases, apart from their placement to putative phage integrases (as the CAT domain of all TRs belongs to the ‘phage_integrase’ Pfam family). Therefore to test the predictive power of our functional annotation we further aimed to identify if some of the non-annotated TRs from the ICE-related subgroups are integrases of ICEs that were not reported previously. For this we examined the genomic neighborhood of the TRs for the presence of known conjugative machinery proteins: if a single integrase was found in close proximity (+/-100Kb) to known conjugation machinery proteins, then the corresponding region was considered to be a putative ICE (conjugation machinery proteins annotated as in [80,81]). ICEs retrieved from ICEberg database were used for benchmarking. The TRs from all the newly identified elements that were previously missing in the ICEberg database (altogether 59 elements) belonged to either of the five AB-containing (ICE/phage) subgroups as based on their match to the HMM profile library (Supplementary Table 4). Of these 2 belonged to Int_KX_, 3 to Int_CTnDOT_, 23 to Int_Tn916_, 17 to Int_P2_ and 14 to Int_SXT_ subgroup. To verify that recovered integrases indeed belong to the corresponding phylogenetic subgroups we then reconstructed the phylogeny of these ICE TRs and plotted their respective conjugation machinery type and structure (Figure 6). This showed moderate evolutionary conservation of the conjugation machinery between ICEs closely related based on their TR phylogeny (Figure 6), and supported previous report on conjugation machinery exchange events happened between more distantly related ICEs [64]. The identification of new ICEs based on the presence of specific TRs validates our functional annotation of the TR subgroups and the utility of our classification system for an automated function prediction.

**Figure 6.**
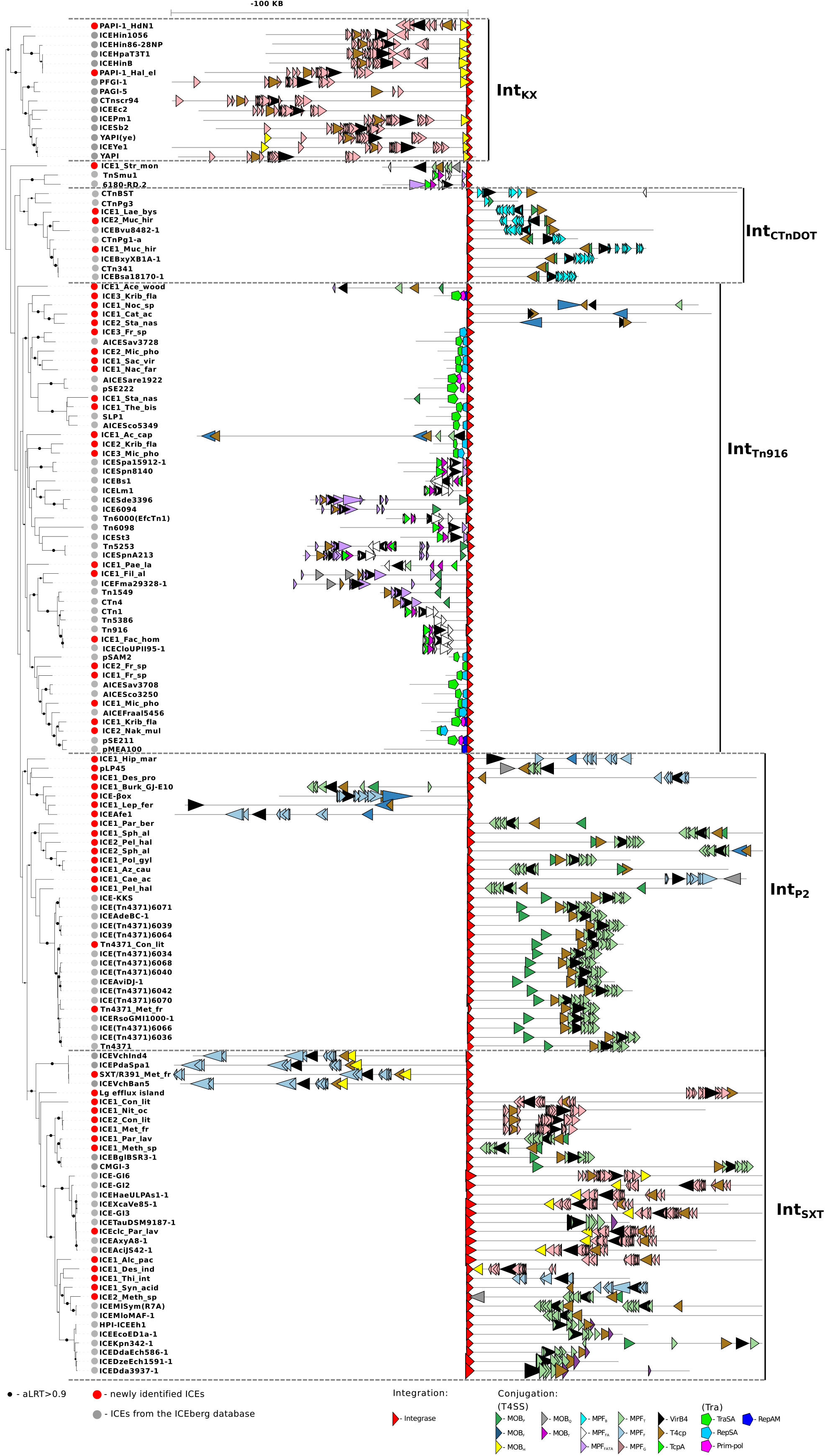
Structural composition of the ICEs. All ICEs are clustered into five subgroups based on their integrase phylogeny. New ICEs that were identified in the present study are marked with the red circles; ICEs from ICEberg database are shown with grey circles. The schemes of ICE architectures are aligned to the integrases (red triangle in the middle). Various types of conjugation machinery proteins are highlighted with different colors.

## DISCUSSION

### Xer tyrosine recombinases are ancient TRs

In the present study, we devise a classification system for bacterial TRs that are related to Xer recombinases. We divide the TRs into two groups and twenty subgroups. Of these, the most abundant and ubiquitous subgroup is Xer, which includes close homologues of the chromosome dimer resolution proteins XerC/D (Figures 1 and S1B). Previous reports indicated a wide distribution of the Xer recombination system in proteobacteria [32]. More recently conserved *dif* sites, the substrates for recombination, were found in 16 bacterial phyla [82]. In agreement, our search revealed a similar distribution pattern for the Xer subgroup TRs (Figure 1B), consistent with the requirement for *dif* sites to coexist with the corresponding Xer proteins in the same genome. Considering that proteobacterial XerC/D proteins were reported to undergo mostly vertical evolution [32], together with our finding that the Xer subgroup has the widest distribution of all TRs studied, it appears that the Xer proteins are the most ancient type of ‘AB-less’ TRs.

### Simple TRs can perform excision and integration of mobile elements

We also report that phages or ICEs can encode simple TRs. These TRs belong to one of two subgroups: the Int_Brujita_ subgroup found in mycophages, and the Int_KX_ subgroup that includes TRs of gammaproteobacterial ICEs (Supplementary tables 1 and 2). In addition, some isolated examples, such as the TnPZ XerT integrase and a putative integrase of ICEBmu17616-1, belong to the Xer and RitB subgroups of simple TRs, respectively. For several of these TRs (e.g. from the Brujita phage, from the Int_KX_ subgroup and XerT), their requirement for excision of the elements has also already been demonstrated [35,43,83,84]. Clustering of these TRs with the Xer subgroup suggests that these phages and ICEs might have captured an ancient Xer from their host genome. However, reverse transition from a mobile element-encoded recombinase to chromosome dimer resolution protein also seems possible. For example, XerH and XerS, functional homologues of the XerC/D proteins in epsilonproteobacteria and in Lactococci respectively, are phylogenetically fairly distantly related to XerC/D [32] and may have been acquired via mobile element transfer [3].

### From simple recombinases to arm-binding domain-carrying integrases

Although some phages and ICEs use simple TRs for their transfer, most of them possess AB domain-containing integrases (27 phages with simple TRs versus 410 with AB-carrying TRs, Supplementary Table 2; and 12 ICEs with simple TRs, 175 with AB-containing TRs and 36 ICEs having multiple types of the TRs, Supplementary Table 1). These integrases belong to one of five subgroups: Int_CTnDOT_, Int_P2_, Int_SXT_, Int_BPP-1_ or Int_Tn916_. In addition, the AB-containing TR group includes the λ phage integrase, which did not cluster with other sequences into a subgroup in our analysis, and the Int_Des_ subgroup, members of which have not yet been functionally annotated. In all these TRs, the N-terminal AB features a similar secondary structure (Figure 5), suggesting a common evolutionary origin for these integrases and their ABs. Moreover, although the AB domain is clearly less conserved than CAT, sequence similarity can be detected between ABs from different subgroups. First of all, the AB Pfam domains from all subgroups belong to the same clan (CL0081), which suggests that they are evolutionarily related. In many cases, however, the AB domains cannot be predicted using Pfam annotations (Figure S1). Nevertheless, the sequence logos created using information from all UniProt reference proteomes proteins show a significant conservation of the N-terminal domain sequences in most of the proteins within each subgroup (Figure 5); and logos of AB domains from different individual TR subgroups reveal marked similarities. These include: a conserved aromatic residue in the second beta strand (in the corresponding third strand in the Int_SXT_ subgroup); a positively charged third beta strand (fourth strand in the Int_SXT_ TRs); a conserved glycine between the last beta strand and C-terminal alpha helix; and a conserved alanine in the C-terminal alpha helix. The similarity between the ABs of various TRs is also supported by structural studies, which revealed a similar core fold (with a specific addition of an alpha-helix and a beta-strand in the Int_SXT_ subgroup TRs) [29,30,85]. The ABs within the Int_CTnDOT_ subgroup, however, seem to be more diverged. Although a similar secondary structure to other ABs was previously predicted for CTnDOT integrase ABs [54], we could not find the conserved sequence patterns we detected for other ABs in the Int_CTnDOT_ ABs (Figure 3), suggesting a distinct origin for the ABs in this group. The functional role of the widespread AB domain may be inferred from its function in the λ phage integrase and biochemical characteristics of the arm-less TRs. The λ integrase AB domain binds to subterminal arm sites within the phage attachment sites and plays an essential role [86] by regulating the directionality of recombination [87,88]. Specifically, the differential spatial arrangement of the arm sites relative to the core recombination sites during excision and integration helps guide λ integrase-mediated recombination towards phage excision or integration [87]. The presence of an AB in most of the ICEs and phage integrases implies that these elements may generally benefit from the action of an AB domain for the regulation of their integration and excision. However, the cases of the Int_KX_ and Int_Brujita_ AB-free TRs show that phage and ICEs mobilization can also be done without AB. These elements may have evolved some other regulatory pathways or may represent an earlier step in evolution of this class of mobile elements [43] with less intricate regulatory features.

## CONCLUSION

Our study provides a comprehensive classification system for Xer-related TRs. For each TR subgroup, we produced an HMM profile and describe characteristic features of its members. In addition, we show that the presence of an additional DNA binding AB domain is characteristic for most phage and ICE integrases that are phylogenetically distinct from simple TRs of non-conjugative transposable elements and from essential bacterial recombinases. Thus, we suggest that sequence homology may be used for functional annotation of the TRs and provide first results of such an annotation, resulting in identification of 59 new putative ICEs. We also propose that phage and ICE integrases originated from Xer homologues by acquisition of an arm-binding domain, since *i*) Xer proteins seem to be the most ancient TR proteins; *ii*) Xer is required for the movement of some mobile elements; *iii*) simple TRs are self-encoded by some mobile elements, perhaps representing an early step in the evolution of the phage and ICE integrases; *iv*) AB-containing TRs, are more frequently represented on intricately regulated mobile elements, perhaps providing a distinct proliferative advantage; and *v*) ABs from diverse TRs share similar structure and AB-containing TRs form a monophyletic clade.

Thus, our study of the diversity, conservation and evolution of the TRs provides a useful framework for future annotation of TRs in newly sequenced genomes. We further expect that our HMM profiles will help improve automated functional annotation of TRs in whole genome sequences and even in metagenomes, where sequences of only short reads are available. The presented sequence logos may also help guide future experimental studies by facilitating functional classification of conserved residues for mutational analysis.

## METHODS

### Identification and phylogenetic analysis of tyrosine recombinases from bacterial genomes

As the first step, we used the XerD amino acid sequence (Genbank accession number CDO12527) as a query in a Jackhammer iterative search against the UniProt reference proteomes database with the e-value threshold parameter set to 1e-30 (converged after 37 iterations; accessed 13.01.2016, http://hmmer.janelia.org/search/jackhmmer currently moved to https://www.ebi.ac.uk/Tools/hmmer/search/jackhmmer; [89]). These settings were used to restrict the analysis to Xer-related TRs and avoid inclusion of distantly related sequences that hamper reconstruction of the phylogenetic tree (data not shown). Notably, the applied threshold excluded the Cre recombinase, which does not cluster with any other known bacterial TR and seems to be more related to TRs of eukaryotic retrotransposons [19,90]. Other well-known TRs, including diverse bacterial or phage TRs for which crystal structures are available (XerD, XerH, integrases of phages λ, HP1 and P2 and integron integrase) and most ICE-encoded TRs from the ICEberg database were included in the study. 187 out of 191 (∼98%) TRs of single TR-containing ICEs (to avoid inclusion of TRs unrelated to integrase function) in ICEberg passed the threshold and were annotated in our study (Supplementary Table 1). All the sequences that passed the threshold (altogether 9,909 proteins) were then clustered with a 40% identity threshold using CD-HIT [91] and the representatives of clusters with size more than one sequence were used for further analysis. This was implemented to exclude singletons and rare sequences with truncations. Then, altogether 866 sequences were aligned using the E-INS-I method from the MAFFT software, recommended for sequences where several conserved motifs are embedded in long unalignable regions [92]. In the alignment, columns containing more than 80% of gaps and the N-terminal region of the alignment (corresponding to AB) were removed. The final alignment is available in Supplementary materials. The phylogenetic tree was constructed using the PhyML package [93] with LG+I+G model of protein evolution and evaluated by ProtTest [94]. In the course of the reconstruction, we built 1000 trees using both NNI and SPR moves as topology search and random tree as starting trees. A tree with the maximum likelihood value was used as the reconstruction of the TR phylogeny (Figure 1A). The branch support was evaluated with aBayes [95]. We further defined subgroups as clades that *i*) include more than four sequences, *ii*) have support value of more than 0.98 and *iii*) exhibit similar domain composition of its members (i.e presence/absence of distinct Pfam domains, [96]). The tree with the domain composition retrieved for each of the sequences from the tree is available as Figure S1 and at http://itol.embl.de/shared/gera for inspection.

The phylogenetic trees for Int_SXT_ and Int_Tn916_ subgroups were produced in the same way as for the general tyrosine recombinases tree. The sequences were mined from the ICEberg database [60] and the UniProtKB phage section, using HMM profiles specific for the subgroups (see section below).

### Annotation of tyrosine recombinase putative function, domain organization, conservation and host organism

In order to annotate all TRs from the UniProt reference proteomes, we created HMM profiles for each of the subgroups. For that, we retrieved the complete sequences of the proteins forming the group and reran the MAFFT alignment of the proteins with the same settings as described above. Then we built an HMM profile for this alignment using HMMER3 [97] and set the gathering threshold for the profile as the lowest score that was produced by the sequences used to build the group. The collection of the 20 HMM profiles specific to each subgroup is available in Supplementary Materials. For each subgroup, its coverage against the Pfam ‘phage_integrase’ family was evaluated using a custom Perl script (Figure S2).

To analyze TR distribution, we mapped all proteins that were retrieved in the UniProt reference proteomes search to their genomic regions and built a plot of the distribution of TRs from each subgroup in bacteria, archaea and viruses (Figure 1B, Supplementary Table 2), visualized using the iTOL webserver [98]. In addition, to shed some more light on the abundance of the TRs in bacterial genomes we extracted fifty of the most abundant TRs found in the sequenced genomes and annotated their distribution in bacteria, classification and putative function (Figure 1C, Supplementary Table 3).

To visualize the similarities and differences between TRs from the different groups and subgroups we created logos representing the sequences from each of the subgroups (Figures 2-5). For that we used the protein alignments, containing all hits produced by the HMM profile-based search against the UniProt reference proteomes database in the previous step, and visualize them using the Skylign webserver [99]. Logo for the λ integrase-related TRs, which failed to form a subgroup as they have only two members of the clade in the tree (Figure S1), was created similarly with only two sequences used for the corresponding HMM profile reconstruction. Secondary structure predictions were performed using Jpred 4 [100].

To retrieve functional annotation of the proteins we run the profile HMM against databases of the proteins with known functions. These included Protein Data Bank (PDB, www.rcsb.org; [101]), ICEberg (Supplementary Table 1; http://db-mml.sjtu.edu.cn/cgi-bin/ICEberg/; [60]) and ACLAME (http://aclame.ulb.ac.be; [102]). In addition, the homologues of the identified TRs were identified using BLAST searches available from ISFinder (database on the insertion sequences, www-is.biotoul.fr; [40]), INTEGRALL (integron databse, http://integrall.bio.ua.pt, [103]) and NCBI CDD ([20]).

### Annotation of new putative ICEs

To predict new ICEs we, first, expand the genomic regions 100 Kb upstream and downstream of the TRs. These regions then were checked for the presence of conjugation machinery proteins using a custom Perl script, which is available from BitBucket repository (https://bitbucket.org/bateman-group/ice_scripts/overview).

Although some ICEs may be up to 600 Kb in length, we chose to cover only +/-100Kb region as a good compromise between the sensitivity of the search and the computational time. All of the full-length ICEs present in the ICEberg database were confirmed using this approach, providing a benchmark for the method. The conjugation machinery proteins were identified using hmmscan search against HMM profiles available from CONJdb [80,81] with an e-value set to 1e-20. All ICEs identified in the study are presented in Supplementary Table 4 and will be submitted to the ICEberg database.

## DECLARATIONS

### Availability of data and materials

All data generated or analysed during this study are included in this published article and its supplementary information files or are available from the corresponding author on request.

### Authors' contributions

GS performed the study. GS, OB and AB designed the study and wrote the manuscript. All authors read and approved the final manuscript.

## Acknowledgements

The authors thank Dr. Michael Chandler for reading and commenting on the manuscript.

