## Supplementary material for "Sequence analysis allows functional annotation of tyrosine recombinases in prokaryotic genomes": Figure S

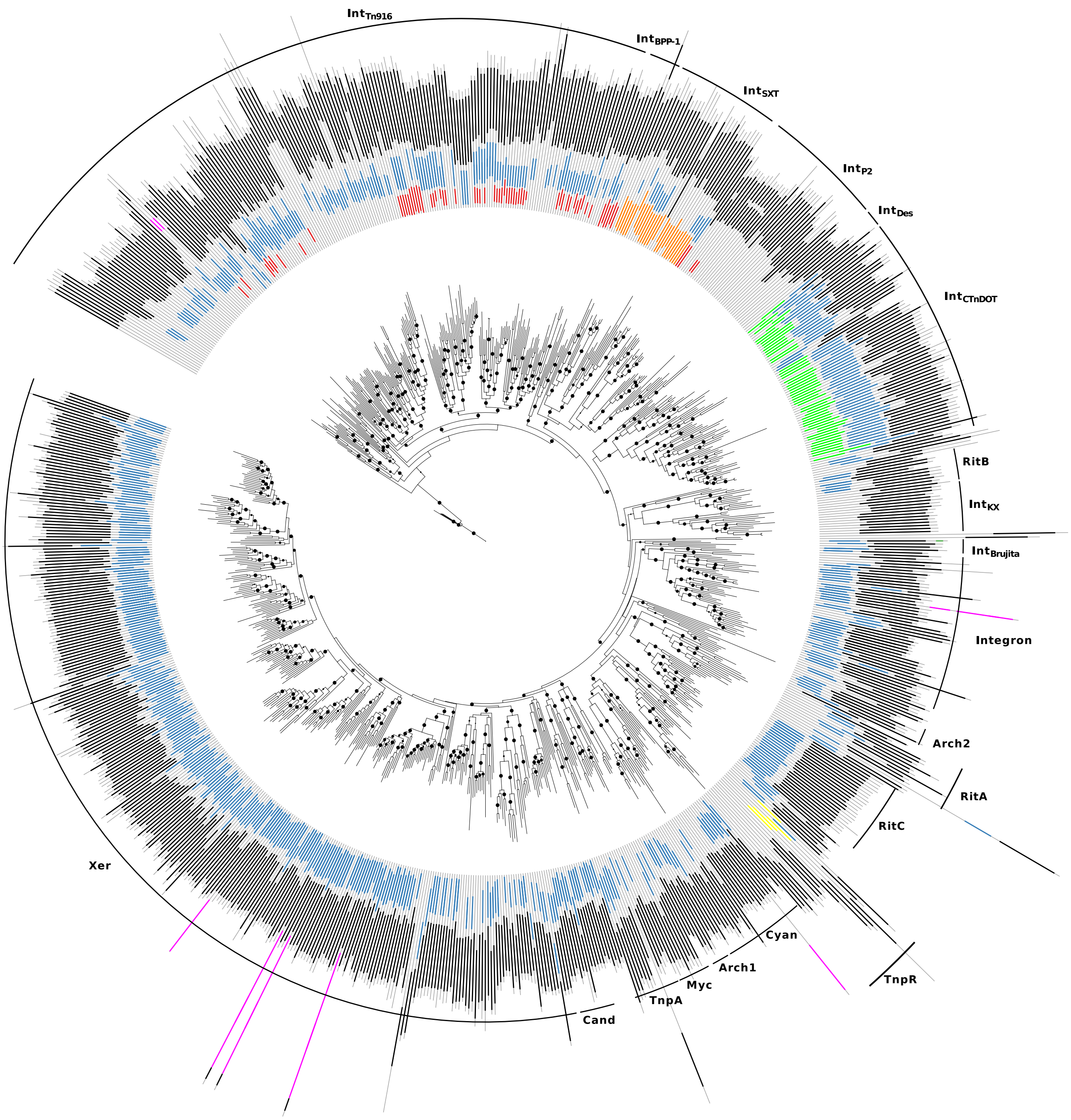


**Protein domains:**

- Catalytic core domain: phage_integrase (PF00589)
- Core binding domain: SAM_1, SAM_2, SAM_4 or SAM_5 (all CL0469)
- DUF3701 (PF12482 )
- DUF3596 or Integrase_AP2  (both CL0081 )
- DUF4102 (PF13356 )
- Arm-DNA-bind_5 (PF17293)
- Other domains

**Supplementary Figure 1.** Maximum likelihood phylogenetic tree of tyrosine recombinases. Statistical support was evaluated by aBayes and values higher than 0.98 are shown as black circles at the corresponding nodes (the larger the circle the higher the value). Subgroups of the TRs are marked outside the tree. The structures of the individual proteins are shown with colored bars representing the predicted domains (color code is shown at the bottom of the figure).

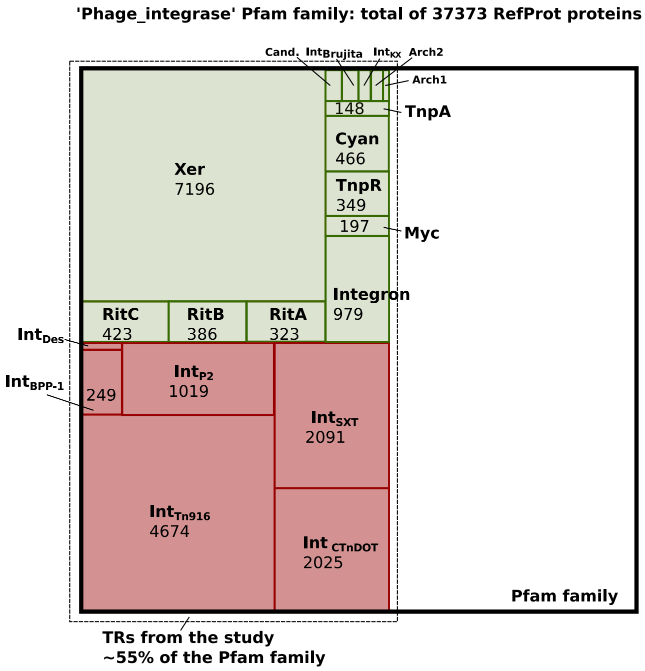


**Supplementary Figure 2.** The coverage of the HMM profiles produced in the study against the Pfam 'phage_integrase' family (Pfam: PF00589). Each square represents the number of hits that were obtained by running a subgroup HMM profile against all the sequences from 'phage_integrase' Pfam family alignment. The exact hit number for the subgroups with more than one hundred hits is shown inside each square.

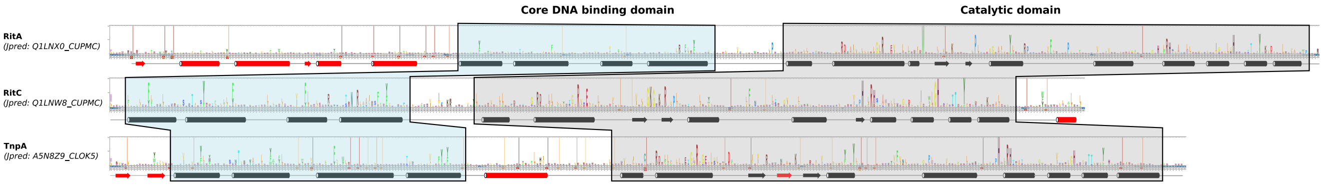


**Supplementary Figure 3.** Full sequence logos of the tyrosine recombinases from RitA, RitC and TnpA subgroups. Different residues are highlighted with different color. Secondary structures were predicted using Jpred and are shown at the bottom of the sequences logos. Secondary structure elements that are specific for each subgroup and missing in other TRs are highlighted in red.

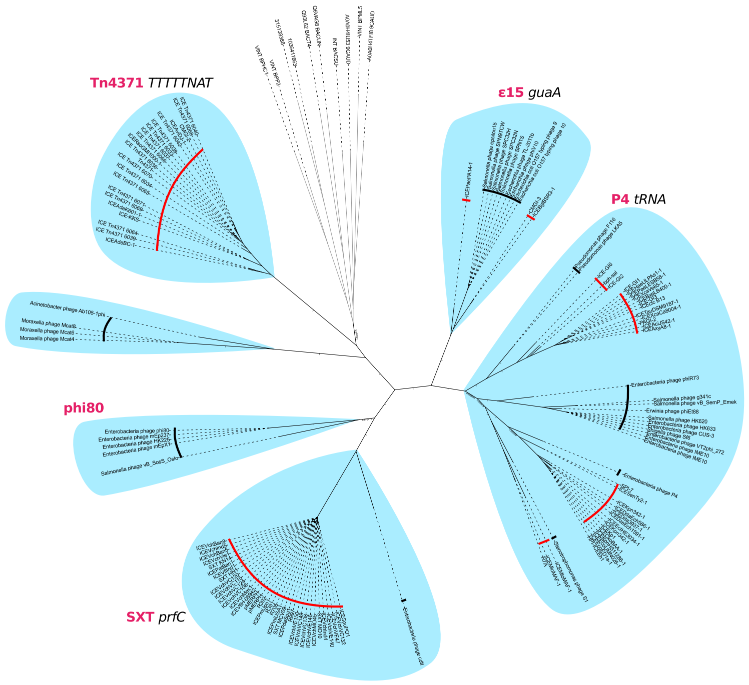


**Supplementary Figure 4.** Maximum-likelihood tree of integrases of ICEs and phages from the IntSXT subgroup. ICE and phage integrases are highlighted in red and black respectively.

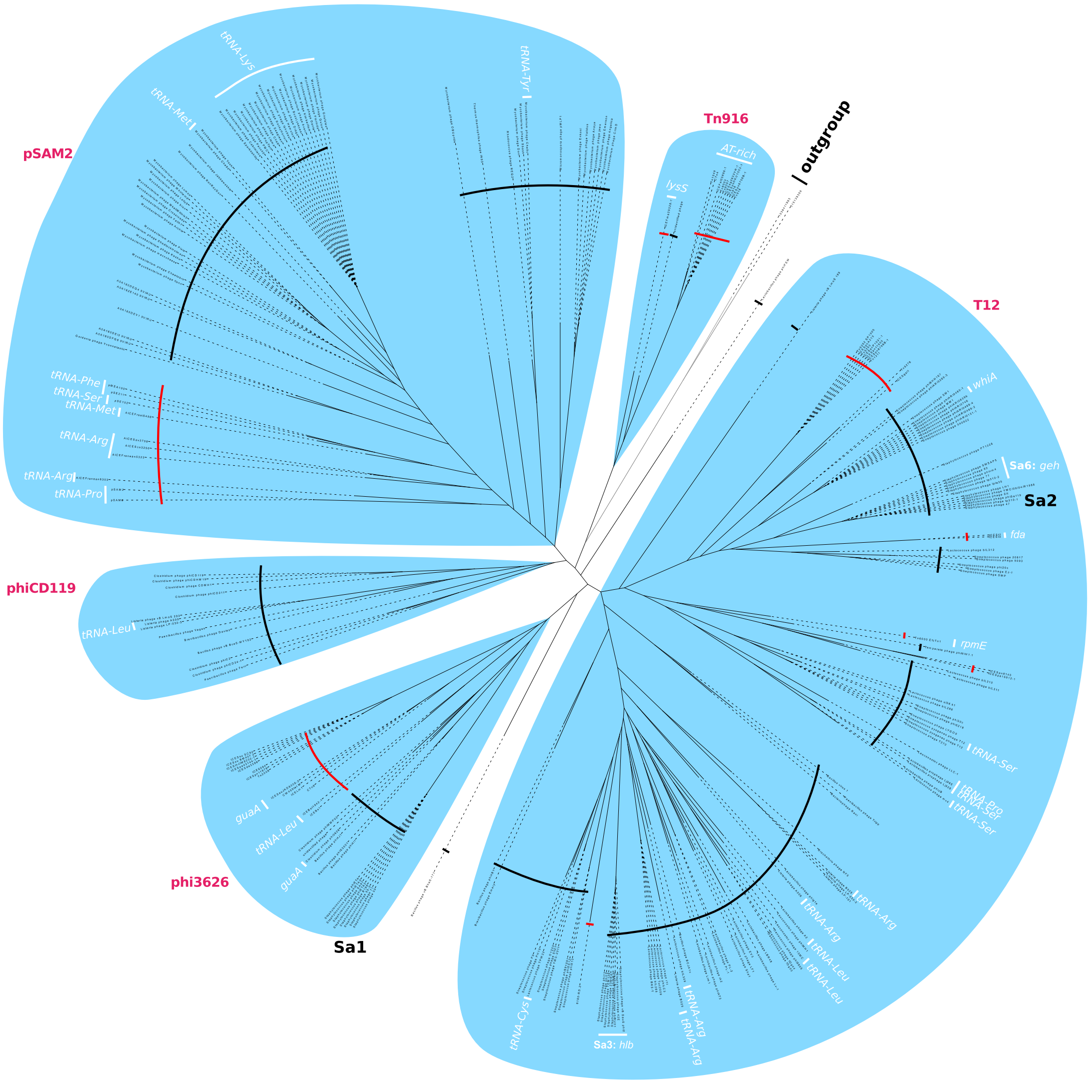


**Supplementary Figure 5.** Maximum-likelihood tree of integrases of ICEs and phages from the IntTn916 subgroup. ICE and phage integrases are highlighted in red and black respectively**.**
